# The effects of post-divergence gene flow on simultaneous divergence time (SDT) testing

**DOI:** 10.64898/2025.12.01.691639

**Authors:** Michael A. Tofflemire, Kevin J. Burns, Jeet Sukumaran

## Abstract

Estimating simultaneous divergence across multiple taxonomic groups from genetic data is a common approach in phylogeographical studies, providing insight into the historical and evolutionary factors shaping species diversification. However, the robustness of divergence time estimates across multiple co-distributed species or population pairs in the face of post-divergence gene flow is often left unaddressed. Here, we use a simulation-based approach to test the robustness of estimating simultaneous divergence within a full-likelihood Bayesian framework when the model assumptions of no gene flow are violated. We generated simulated datasets of multiple population pairs with varying migration rates and estimate shared divergence times with the software package Ecoevolity, comparing them with estimates from population pairs experiencing no migration. Our goal was to identify the threshold at which migration rates begin to bias divergence time estimates, providing insight into the robustness of full-likelihood Bayesian methods under more complex demographic scenarios. We found that simultaneous divergence is incorrectly supported across a broad range of parameters space when assumptions are violated. We suggest that future empirical studies that use simultaneous divergence testing explicitly test for gene flow, or at the very least, explicitly consider the presence, absence, and implications of gene flow in their systems as part of their investigations. We assert that this result—biased support for simultaneous divergence—is just a specific example of a general behavior when the statistical assumptions of the model are not met or there is insufficient power given the data. We additionally recommend that future studies reorient their research framework to focus on trying to demonstrate non-simultaneous divergence in systems. As such, support for simultaneous divergence between groups should be interpreted as failure to resolve non-simultaneous divergence rather than evidence of true simultaneous divergence.

## Introduction

Estimating simultaneous divergence across multiple taxonomic groups from genetic data is a common approach in phylogeographical studies, providing insight into the historical and evolutionary factors shaping species diversification (Hickerson et al., 2006; Huang et al., 2011). At the community scale, concordance in phylogeographic patterns across the landscape provides strong evidence for major historical and geological processes driving regional patterns of biodiversity (Avise, 2000). For example, habitat fragmentation resulting from glaciation events (e.g., Pleistocene glacial cycling) can cause entire communities to fragment within multiple isolated refugia, resulting in shared patterns of spatiotemporal genetic differentiation across multiple co-distributed taxonomic groups (Hewitt, 2000). Conversely, species-specific traits, including habitat preference, can also mediate a taxon’s response to environmental changes, leading, instead, to non-synchronous patterns (Papadopoulou & Knowles, 2015). Thus, using statistical methods to quantify temporally clustered events across multiple co-distributed taxa is essential for testing predictions of community-scale processes.

Analyses aimed at inferring the degree to which divergence patterns are shared across different groups of species and population pairs– based on shared divergence times taken as support for population genetic evolution being structured by shared geographical or other external events – are commonly referred to as “simultaneous divergence time testing” in the literature (Hickerson et al., 2006). While details might vary, with inference being implemented in both approximate Bayesian computation (ABC; Hickerson & Stahl, 2006) as well as full-likelihood Bayesian inferential frameworks (Oaks et al., 2020), all of these analyses are based on optimizing the number of time-separated coalescent-structuring events required to explain variation in genetic data sampled from multiple pairs populations of populations assumed to have diverged from some common ancestor shared exclusively between them. These analyses aim at selecting among different partitions of the set of species groups into clusters of shared “co-diverging” groups, with model preference based on either probability or summary statistic distance functions in full-likelihood or approximate-likelihood Bayesian approaches, respectively. These methods have had demonstrated success in revealing insight across various systems that would not have otherwise been possible due to lack of reliable fossils for independent time-calibrated divergence time estimation to correlate phylogenetic branching events with hypothesized geological or demographic events (Bermingham & Moritz, 1998).

The coalescent and population genetic theory that motivate the probability functions (e.g., in full likelihood inference analyses) or the summary statistics (e.g., in ABC analyses) make many simplifying assumptions about the systems and processes that generated the data, some of which may be violated when applied to real-world data. The assumptions of the multi-population coalescent models underlying the analyses for testing simultaneous divergence time across population pairs are well known and reported. For example, the most basic models include constraints on population sizes (e.g., no population size differences between populations within or across species, and no changes in population size between events), the mutation process (e.g., shared and constant mutation rates, with sufficiently high mutation rate given the time constraints to result in sufficient genetic variation to inform the model), migration (e.g., no migration between any of the descendent populations following a split), and within-population lineage structuring (e.g., ancestral or daughter populations are completely panmictic, with no neighborhood or isolation by distance effects). More advanced models relax these assumptions by, for example, explicitly incorporating these processes at a population level with parameter that is free to vary between populations to explain some of the genetic variation. However, beyond the fact that some subsets of parameters might be confounded, rendering models non-identifiable if all are free to vary, the more free parameters allowed in a model, the lower the statistical power. As such it would seem to be good practice for anyone applying these approaches to first test their system for evidence of violation of these various constraints, and then introduce the required model parameter in piecemeal, as needed, so as maximize statistical power while minimizing complexity.

Despite, we found that of 57 published studies reviewed – 23 of which used a full likelihood Bayesian approach for simultaneous divergence testing –only 6 tested for violation of the assumptions of no gene flow, for example, and only 10 discussed or acknowledge this as a potential issue without explicitly testing for them or otherwise accounting for them. As such, the robustness of divergence time estimates across multiple species or population pairs in the face of this violation of no gene flow is often left unaddressed in studies. It would be unfair to place too much burden of omission on these empirical studies however, for it is not always straight-forward to determine if some assumptions have been violated statistically, and more importantly, there has not yet been any theoretical analysis or study that characterizes any of these issues in any quantitative detail to provide the fundamental understanding required so that an empirical investigator can make informed decisions about their system by being able to recognize or even predict behavior or rather, “misbehavior”, of these analyses when their underlying assumptions are violated.

Simultaneous divergence testing (henceforth, SDT) has been implemented in both approximate likelihood and full-likelihood Bayesian frameworks. Approximate-likelihood Bayesian frameworks use summary statistic distances between data simulated from competing models and the observed data to approximate the likelihood when sampling from the posterior, this latter typically achieved through by rejection sampling of the simulations. In both approaches, there is no theoretical issue with taking migration into account. In the case of the full-likelihood approaches, expressions are available for post-divergence migration, while in the approximate-likelihood approaches, post-divergence gene flow is already available as an optional parameter in the underlying simulator model of all implementations of the analyses.

Despite this, no published study has opted to consider post-divergence gene flow, and there has been no published investigation on the impact of post divergence gene flow. Here, we generate simulated datasets of multiple population pairs with parameters representing varying levels of migration rates and estimate shared divergence times in the program Ecoevolity, taking advantage of the greater statistical power afforded by a full-likelihood Bayesian approach over ABC, and comparing them with estimates from population pairs experiencing no migration to: (1) Quantify the relative threshold at which migration will have an impact on inference of shared divergence time patterns; (2) Assess whether migration primarily affects inferential reliability (as measured by confidence interval coverage and model selection accuracy) or estimation precision (as measured by root mean squared error); (3) Determine if observed errors represent model misspecification bias (systematic failure to capture true parameters in confidence intervals) or simply increased parameter uncertainty (reduced power while maintaining calibrated inference), and (4) Evaluate the impact on model selection: whether migration leads to incorrect inference of divergence event patterns (systematic model selection bias) or simply reduces power to distinguish between models.

## Methods

### Simulating population pair nucleotide baseline datasets without gene flow

Simulated datasets were generated using the msprime and tskit software packages under a range of evolutionary models (Baumdicker et al., 2022; Kelleher et al., 2016; Nelson et al., 2020). For each dataset, we simulated DNA sequences for six population pairs. Our baseline simulation parameters included 20 haploid DNA sequences per population pair (10 sequences per population), with each sequence consisting of 1 locus of 1,000 base pairs (bp) and generated with a mutation rate (*µ*) of 1.0 × 10−8 mutations/site/year (Nachman & Crowell, 2000). A generation time of 1 year was used for computational simplicity.

For the baseline demographic model, each population pair diverges from a common ancestor *t* generations ago with an effective population size (*N_e_*) of 10,000 individuals. Following the split, each population retains an *N_e_* of 10,000. No gene flow occurs between populations after the split (i.e., strict isolation model). Within each population, there is no further substructure, and no recombination or natural selection occurring. Sequences were simulated under a coalescent model (4*N_e_*µ).

We simulated each dataset at various configurations of the six population pairs by varying the timing of divergence (*t*) of each population pair (where *t* represents generation time) or the time in between splitting events (*Δt*) between each population pair (see Figure 1). We simulated a total of three configurations: (1) all six population pairs share the same divergence time, where number of shared divergence events (*N*_div_) equals one; (2) all six population pairs have unique divergence times, where *N*_div_= 6; and (3) pairs grouped in threes with the same divergence time, where *N*_div_= 3 (see Figure 1).

**Figure 1.**
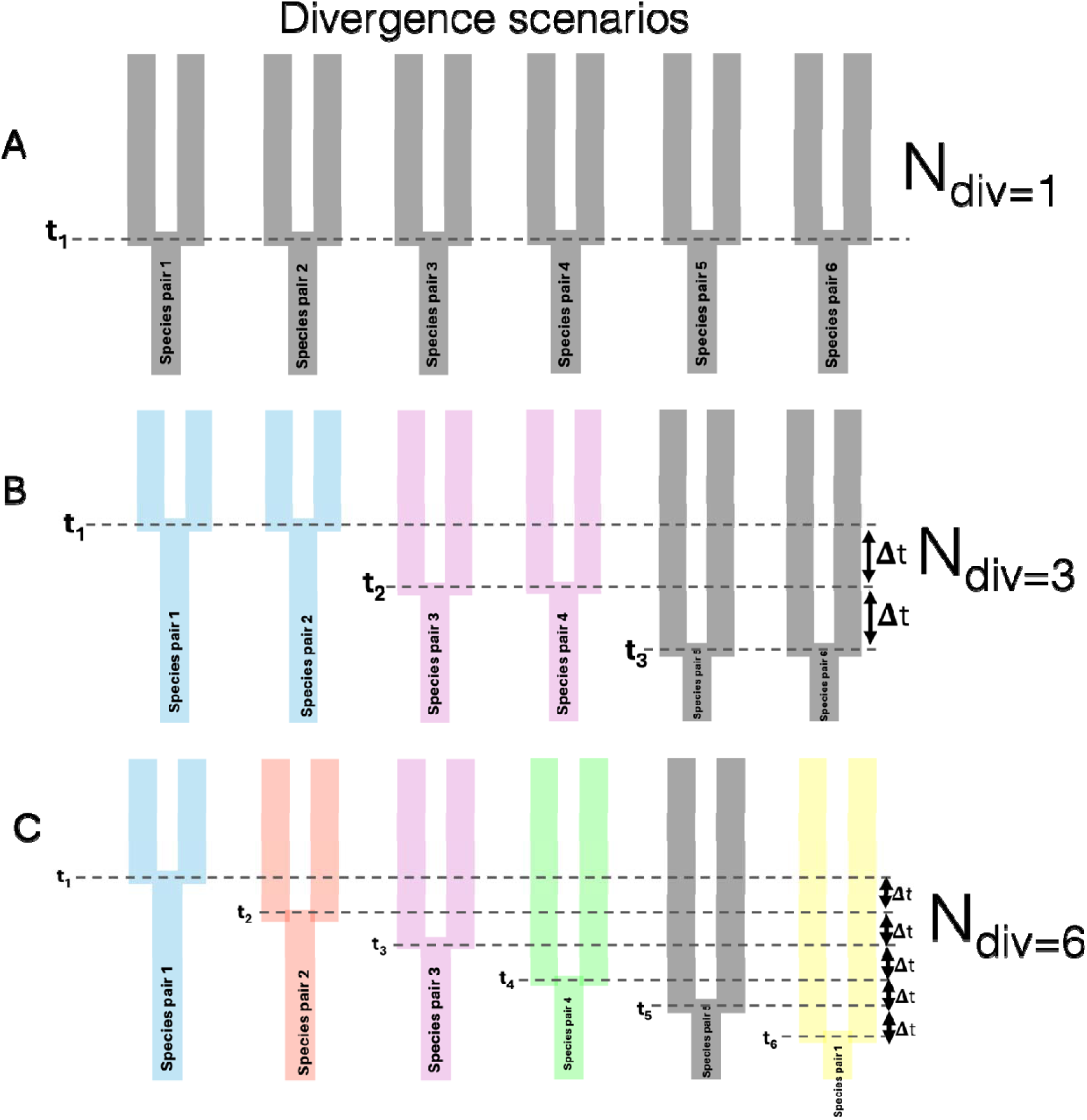
Illustration of three configuration models based on six population pairs. (a) The first configuration represents a single divergence event shared across the six population pairs. (b) Three groups of shared divergence events. (c) Six independent divergence events across the six population pairs. Dashed lines represent time of divergence event (t). represents time in between splitting events.

For the *N*_div_ = 1 configuration, where all six population pairs undergo simultaneous divergence, we simulated 5 separate datasets where all population pairs share divergence times of 10,000, 20,000, 40,000, 80,000, and 160,000 generations ago. Since *N_e_* was 10,000 for all populations, divergence times above correspond to 1*N_e_* (more recent splits), 2*N_e_*, 4*N_e_*, 8*N_e_*, and 16*N_e_* (more distant splits) generations ago, where *N_e_* represents the ancestral effective population size.

For the *N*_div_ = 6 configuration, we began with a baseline divergence of t = 10,000 generations for the first population pair. For each subsequent simulated pair, we increased the time separation between pairs by a fixed amount (Δt), which represents the interval between splitting events. For each simulation, Δt was set to 10,000, 20,000, 40,000, 80,000, or 160,000 generations in the positive direction starting from 10,000. These correspond with a Δt of 1*N_e_*, 2*N_e_*, 4*N_e_*, 8*N_e_*, and 16*N_e_* generations. For example, in scenarios where Δt was set to 10,000, divergence times for the six population pairs –– A|B, C|D, E|F, G|H, I|J, and K|L –– were set to 10,000, 20,000, 30,000, 40,000, 50,000, and 60,000 generations, respectively. These same settings were applied to the third configuration, producing 15 unique datasets across all configurations without gene flow.

### Simulating population pair nucleotide datasets with gene flow

The same configurations and demographic scenarios described above were simulated a second time except with varying levels of gene flow occurring between population pairs to see the effects migration has on estimating divergence times across species. Gene flow (i.e., migration) was varied at multiple rates including a high migration rate (*m)* of 1.0 × 10−2 individuals/generation, and low migration rate of 1 × 10−7 individuals/generation.

### Running Ecoevolity and assessing performance for each model

For each simulated dataset, we used Ecoevolity to estimate the number of shared divergence events and the timing of the most recent common ancestor (MRCA) for each divergence event across all six population pairs. We then compared these estimates to the true values within a statistical framework to assess the threshold at which we could examine non-synchronous divergence. Briefly, Ecoevolity uses a full-likelihood Bayesian approach to quantify the most likely number of shared divergence events. Key assumptions of the model include no gene flow, orthologous genetic markers, biallelic sites, and unlinked markers. To prepare our simulated datasets, sequences were converted to nexus format and combined into a single file containing all population pairs. For each dataset, we ran 30 replicates, compiling summary statistics that included the initial and estimated parameters for probability of shared events and divergence times for each population pair. For each replicate, we used the Ecoevolity default configuration, with a Dirichlet process, MCMC chain length of 1,000,000, and a sample frequency of 100. Convergence was checked for each output to ensure ESS > 200, and results were visualized using R packages.

Performance was quantified using two primary metrics: (1) root mean squared error (RMSE) of divergence time estimates relative to true simulated values, and (2) coverage probability at the 95% confidence level (the proportion of replicates in which the true divergence time fell within the 95% credible interval). We assessed performance across three divergence scenarios—simultaneous divergence across all six population pairs, three pairs of synchronized divergences, and six independent divergences —each simulated at five divergence ages (1*N_e_*, 2*N_e_*, 4*N_e_*, 8*N_e_*, 16*N_e_* generations) and three post-divergence migration rates (0, 1 × 10−7, 1 × 10−2).

## Results

### Baseline cases: no gene flow

Simultaneous divergence in the absence of post-divergence gene flow across all six population pairs showed perfect coverage probabilities (p(t ∈ CI) = 1.00) across all divergence ages (Figure 1a, top row). RMSE values were minimal, ranging from 2.13 × 10−4 at 1*N_e_* to 5.80 × 10−4 at 16*N_e_* generations, showing a modest increase with divergence age. Grouped divergences for three pairs of shared events (Figure 1a, middle row) showed reduced coverage probabilities compared to the fully simultaneous case. Coverage ranged from 0.667 at shallow divergences (2*N_e_*, 4*N_e_*) to 1.00 at the extremes (1*N_e_* and 16*N_e_* generations), with p(tCI) = 0.833 at 8*N_e_*. RMSE values increased more substantially with age, from 3.54 × 10−4 at 1*N_e_* to 9.45 × 10−4 at 16*N_e_* generations. Finally, the most complex scenario of six independent divergences exhibited the weakest coverage probabilities in the absence of migration (Figure 1a, bottom row). Coverage was lowest at shallow to intermediate divergence times, with p(t ∈ CI) = 0.667 at both 1*N_e_* and 2*N_e_* generations, increasing to 0.833 at 4*N_e_*, 8*N_e_*, and 16*N_e_*. RMSE values were highest among all three scenarios, ranging from 2.91 × 10−4 at 1*N_e_* to 1.74 × 10−3 at 8*N_e_*, with a slight decrease to 1.37 × 10−3 at 16*N_e_*.

We evaluated model selection performance by examining the posterior probabilities assigned to different numbers of divergence events (1-6 events) for no migration (Figure 1b). Model selection under the no migration strongly favored the true single-event model across all divergence ages (Figure 1b, top row). At 1*N_e_* split time separation, probability mass was concentrated almost entirely on 1 event (purple shading shows high probability near probability = 1.0). As divergence age increased, a small amount of probability mass shifted to 2-event models, but the dominant mode remained at 1 event with high probability values. At 16*N_e_*, the distribution showed increased spread with modest probability assigned to 2-3 events, but 1 event remained clearly dominant.

For grouped divergence events of 3 events, the posterior under no migration correctly distributed probability mass across multiple events, with the mode progressively shifting toward higher event numbers as divergence age increased (Figure 1b, middle row). At 1*N_e_*, the dominant mode was at 1-event models, but substantial probability mass appeared for 2-3 events. At 2*N_e_*, the distribution became more diffuse with comparable probability for 1-2 events. By 4*N_e_* and 8*N_e_*, the distribution showed clear bimodality or trimodality with peaks for 1, 2, and 3 events (orange shading indicates moderate probability values). At 16*N_e_*, 2-event models received the highest probability mass, with substantial probability also assigned to 1 and 3 events.

For 6 independent divergence events, the posterior distributions under no migration showed progressive increase in estimated event complexity with divergence age, though consistently underestimating the true six events (Figure1b, bottom row). At 1*N_e_*, the dominant mode was at 1 event. At 2*N_e_*, probability mass shifted to 2-3 events. By 4*N_e_* and 8*N_e_*, the distributions peaked at 3-4 events (substantial pink shading), with 8*N_e_* showing the highest mode at 3 events. At 16*N_e_*, the distribution showed a broad peak centered on 4 events, with probability mass extending from 2-5 events.

### Low migration rate (1×10−7)

Simultaneous divergence scenario at low migration had minimal impact on inference quality in the simultaneous divergence scenario (Figure 2a, top row). Coverage probabilities remained perfect (p(t ∈ CI) = 1.00) across all divergence ages. RMSE values were comparable to the no-migration case, ranging from 1.07 × 10−4 at 1*N_e_* to 5.92 × 10−4 at 16*N_e_* generations. The grouped divergence scenario (Figure 2a, middle row) showed coverage probabilities under low migration that closely paralleled the no-migration case: p(t∈ CI) = 0.833 at 1*N_e_*, 0.667 at 2*N_e_* and 4*N_e_*, 0.833 at 8*N_e_*, and 1.00 at 16*N_e_*. RMSE values were similar to baseline conditions, ranging from 2.45 × 10−4 at 1*N_e_* to 8.64 × 10−4 at 16*N_e_*. Finally, our results for the independent divergence scenario at low migration (Figure 2a, bottom row) produced coverage probabilities of 0.833 at 1*N_e_*, 0.833 at 2*N_e_*, 1.00 at 4*N_e_*, 0.833 at 8*N_e_*, and 0.833 at 16*N_e_* —comparable to or slightly improved from the no-migration baseline. RMSE values ranged from 3.91 × 10−4 at 1*N_e_* to 1.58 × 10−3 at 16*N_e_*, showing similar patterns to the no-migration case.

**Figure 2.**
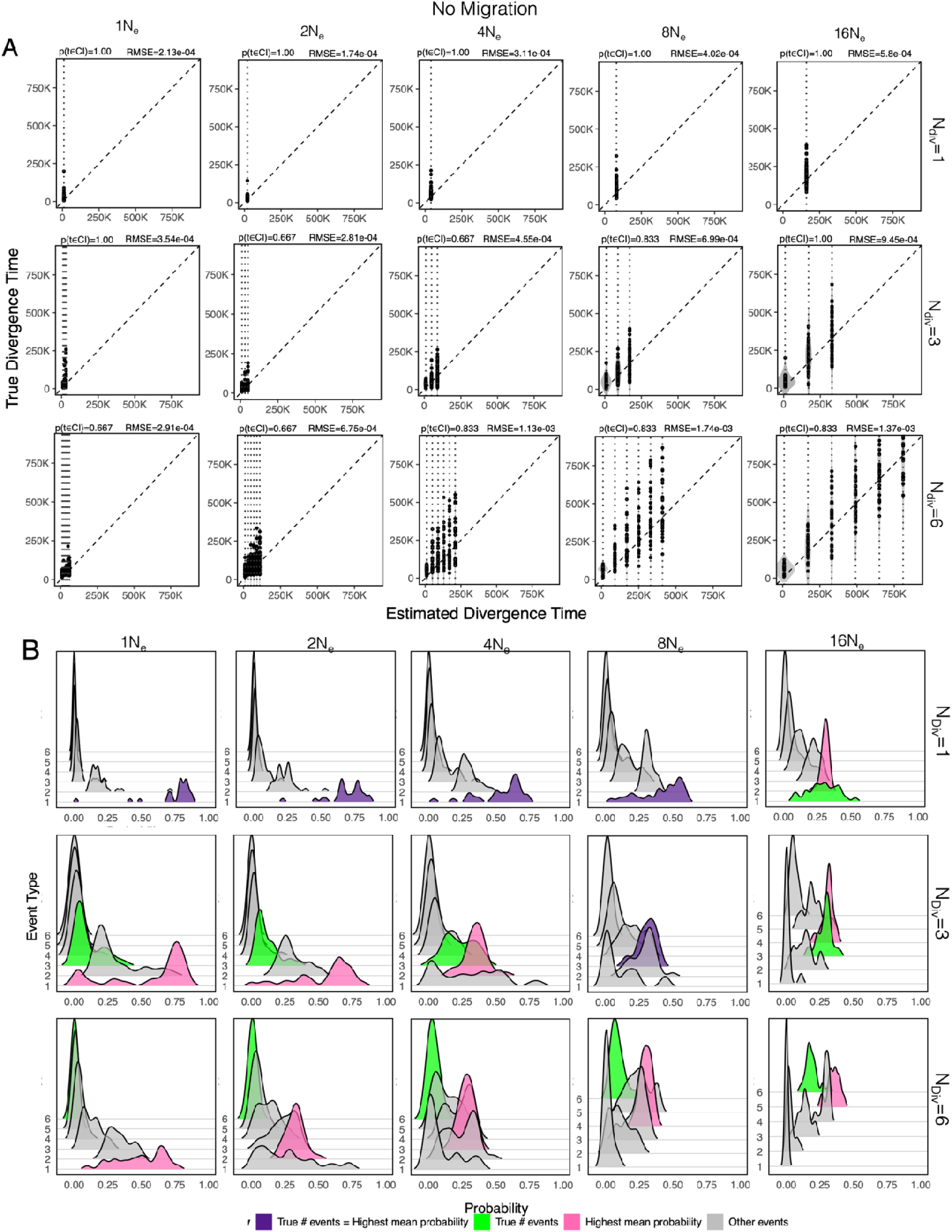
(a) Performance of time estimates for datasets simulated under no migration. Each row corresponds to a particular configuration of the six population pairs (see Figure 1), and each column corresponds to a shared divergence time in generations ago (top row panels) or time separation in-between splitting events (middle and bottom rows) denoted above panel columns as 1*N_e_*, 2*N_e_*, 4*N_e_*, 8*N_e_*, or 16*N_e_* generations, where *N_e_* equals 10,000. Points represent the estimated divergence times in Ecoevolity for each population pair for a single run. Each dataset was run at 30 replicates. The dashed diagonal line represents the one-to-one expectation between the true and estimated divergence times. Metrics for percent coverage of the true value and RMSE are shown above each panel. (b) Posterior probability distributions for the estimated number of shared divergence events. Shading indicates either the event type showing the highest probability (pink), the true probability (green), or when the event with the highest probability equals the true probability (purple).

Posterior distributions under low migration closely resembled the no-migration case (Figure 2b). Strong concentration on 1-event models at shallow divergences, with gradual increase in probability mass for 2-event models at older ages. The distributions maintained the correct identification of a single divergence event as most probable across all conditions. Progressive shift from 1-2 events at shallow divergences to 2-3 events at intermediate and old divergences. At 8*N_e_*, the distribution showed clear support for 2-3 events with pink shading indicating moderate to high probabilities. Posterior distributions under low migration closely paralleled no-migration patterns. Progressive increase from 1 event at 1*N_e_*, to 2-3 events at 2*N_e_* and 4*N_e_*, to 3-4 events at 8*N_e_* and 16*N_e_*. The distributions maintained similar shapes and probability mass distributions to the baseline case, correctly showing increased event complexity with divergence age.

### High migration rate (1×10−2)

Simultaneous divergence scenario at high migration (Figure 3a, top row) resulted in complete failure of confidence interval coverage across all divergence ages (p(t ∈ CI) = 0.00 for all conditions). RMSE values were generally lower than baseline at shallow divergences (7.84 × 10−5 at 1*N_e_*, 1.78 × 10−4 at 2*N_e_*) but increased substantially at older divergences (3.79 × 10−4 at 4*N_e_*, 7.77 × 10−4 at 8*N_e_*, 1.58 × 10−3 at 16*N_e_*). The grouped divergence scenario (Figure 3a, middle row) under high migration also showed complete coverage failure (p(t ∈ CI) = 0.00) across all ages. RMSE values ranged from 1.99×10−4 at 1*N_e_* to 2.13×10−3 at 16*N_e_*, representing the highest error rates observed across all conditions. Notably, RMSE at 1*N_e_* was higher under migration than without, despite the lower coverage. Independent divergences show high migration (Fig. 3a, bottom row) produced zero coverage (p(t ∈ CI) = 0.00) across all divergence ages in the independent divergence scenario. RMSE values ranged from 3.74 × 10−4 at 1*N_e_* to a peak of 2.48 × 10−3 at 8*N_e_*, before decreasing to 1.37 × 10−3 at 16*N_e_*. The 16*N_e_* condition showed the highest RMSE observed in any experimental treatment.

**Figure 3.**
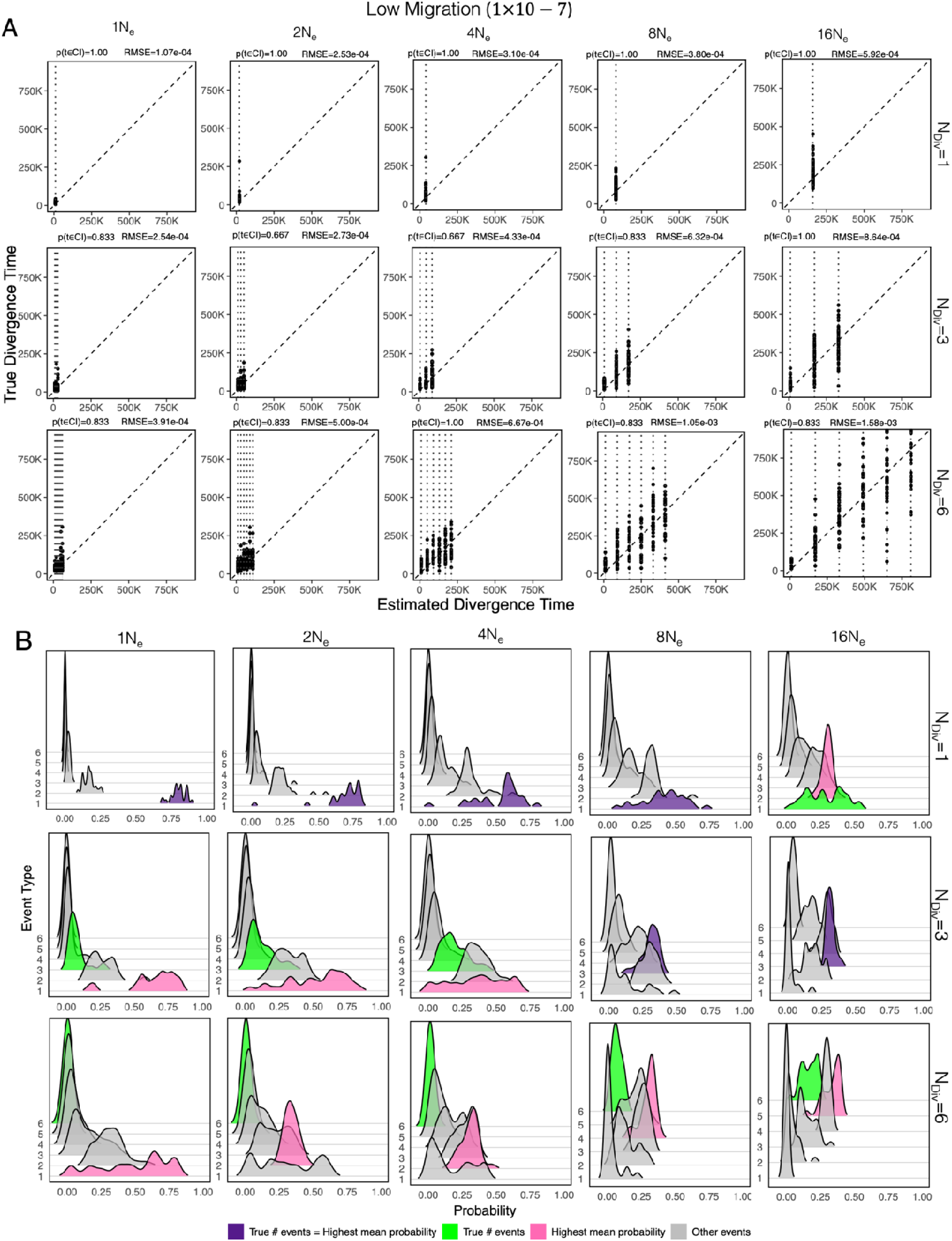
(a) Performance of time estimates for datasets simulated under low migration (1X 10 - 7). Each row corresponds to a particular configuration of the six population pairs (see Figure 1), and each column corresponds to a shared divergence time in generations ago (top row panels) or time separation in-between splitting events (middle and bottom rows) denoted above panel columns as 1*N_e_*, 2*N_e_*, 4*N_e_*, 8*N_e_*, or 16*N_e_* generations, where *N_e_* equals 10,000. Points represent the estimated divergence times in Ecoevolity for each population pair for a single run. Each dataset was run at 30 replicates. The dashed diagonal line represents the one-to-one expectation between the true and estimated divergence times. Metrics for percent coverage of the true value and RMSE are shown above each panel. (b) Posterior probability distributions for the estimated number of shared divergence events. Shading indicates either the event type showing the highest probability (pink), the true probability (green), or when the event with the highest probability equals the true probability (purple).

**Figure 4.**
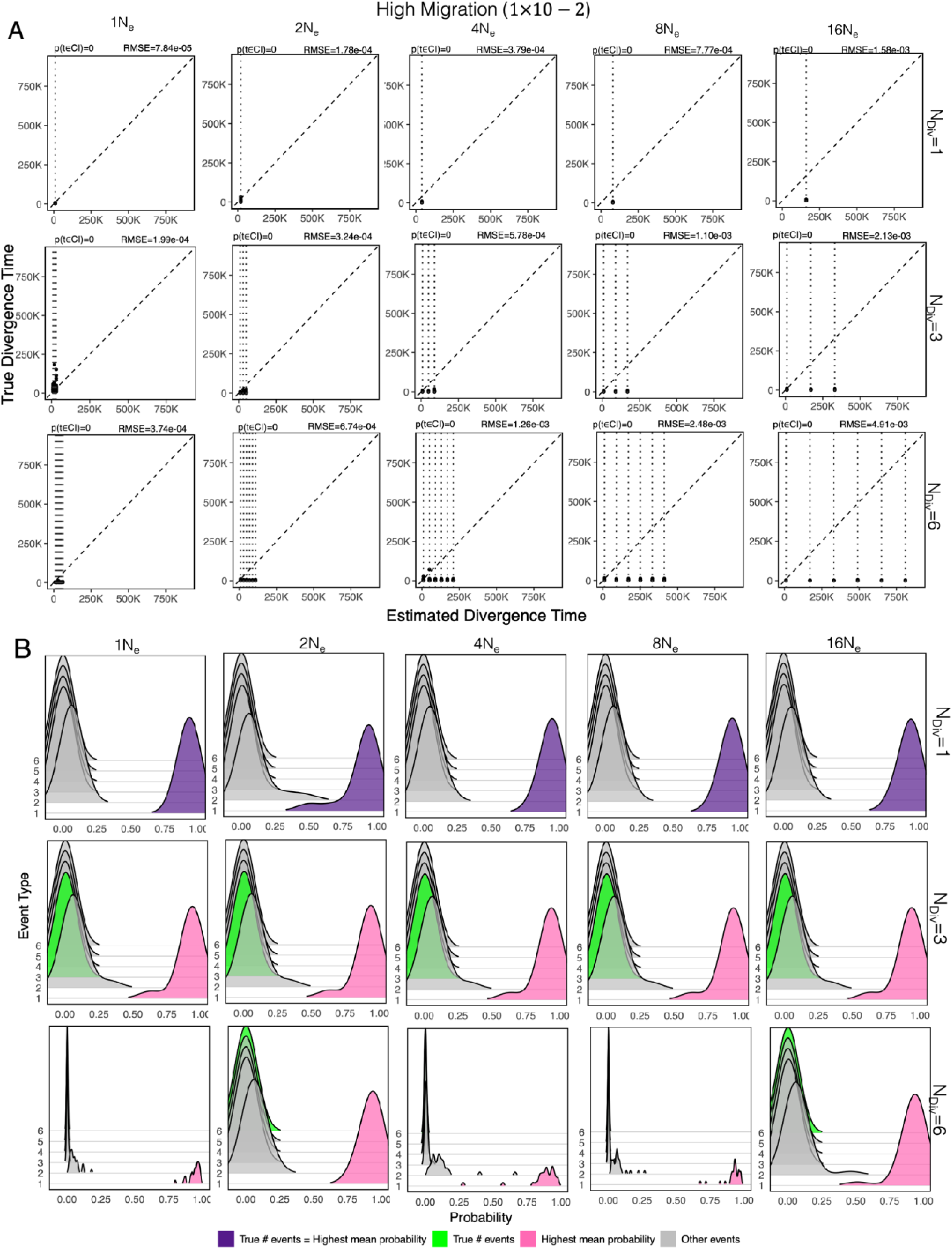
(a) Performance of time estimates for datasets simulated under high migration (1 X 10 - 2). Each row corresponds to a particular configuration of the six population pairs (see Figure 1), and each column corresponds to a shared divergence time in generations ago (top row panels) or time separation in-between splitting events (middle and bottom rows) denoted above panel columns as 1*N_e_*, 2*N_e_*, 4*N_e_*, 8*N_e_*, or 16*N_e_* generations, where *N_e_* equals 10,000. Points represent the estimated divergence times in Ecoevolity for each population pair for a single run. Each dataset was run at 30 replicates. The dashed diagonal line represents the one-to-one expectation between the true and estimated divergence times. Metrics for percent coverage of the true value and RMSE are shown above each panel. (b) Posterior probability distributions for the estimated number of shared divergence events. Shading indicates either the event type showing the highest probability (pink), the true probability (green), or when the event with the highest probability equals the true probability (purple).

Results for model performance under high migration showed the posterior distributions shifting dramatically (Figure 3b). At 1*N_e_* and 2*N_e_*, probability mass became even more concentrated on the 1-event model (very narrow peaks near probability = 1.0). At 4*N_e_*, the distribution began to broaden, with a distinct peak appearing for 2-event models at intermediate probability values. By 8*N_e_* and 16*N_e_*, the distributions showed bimodal patterns with substantial probability mass on both 1-event and 2-event models, with individual peaks at different probability values.

For model performance of the three grouped divergence events, high migration dramatically altered the posterior distributions, strongly biasing them toward fewer events (Figure 3b, middle row). At 1*N_e_* to 16*N_e_*, distributions showed extreme concentration on 1-event models (very narrow peaks at high probability). For all divergence scenarios, the probability mass remained concentrated at higher probability values than the true 3-event model would predict, indicating consistent underestimation of event complexity.

For six independent divergence events, high migration severely biased event number estimation downward (Figure 3b, bottom row). At 1*N_e_*, the distribution showed extreme concentration on 1 event. At 2*N_e_*, probability remained heavily weighted toward 1 event with minimal spread. At 4*N_e_*, 8*N_e_*, and 16*N_e_*, distributions showed consistent peaks at 2-3 events—far below the true value of 6—with probability mass concentrated at intermediate probability values rather than spreading across the full range of possible event numbers.

### Summarization of SDT patterns across all scenarios

The average coverage across all conditions was 0.867 for no migration, 0.889 for low migration, and 0.000 for high migration. Mean RMSE increased with evolutionary complexity: 4.21×10−4 for simultaneous divergences, 6.34×10−4 for grouped divergences, and 1.27 × 10−3 for independent divergences, averaged across all migration rates and ages. Within each scenario, RMSE generally increased with divergence age, though this trend was less consistent under high migration. The ratio of RMSE under high migration to no migration ranged from 0.37 to 5.44 across conditions, with the highest ratios observed in the grouped divergence scenario at older ages.

Finally, high migration consistently shifted posterior probability mass toward simpler models (fewer events). The magnitude of this bias increased with both true model complexity and divergence age. In the simultaneous scenario (true = 1 event), high migration either maintained or slightly increased concentration on 1-event models. In the grouped scenario (true = 3 events), high migration shifted the mode from 2–3 events (baseline) to 1–2 events. In the independent scenario (true = 6 events), high migration reduced the modal estimate from 3–4 events (baseline) to 2–3 events at older divergences. Low migration showed no systematic deviation from no-migration baselines in model selection across any scenario or divergence age, with posterior distributions maintaining appropriate support for the correct level of model complexity.

## Discussion

### Migration threshold and inferential reliability

We assessed how post-divergence migration affects the ability of Ecoevolity to accurately infer shared divergence patterns across three simulation scenarios including simultaneous divergence across all six population pairs, three pairs of synchronized divergences, and completely independent divergences. Each scenario was simulated across a range of divergence ages (1*N_e_*, 2*N_e_*, 4*N_e_*, 8*N_e_*, 16*N_e_* generations) and three migration rates: none (0), low (1 × 10−7), and high (1 × 10–2). Low migration rates (1 × 10−7) had negligible impact on inference quality, with 95% confidence interval (CI) coverage averaging 88% across all conditions—statistically indistinguishable from the no-migration baseline (89% coverage). In stark contrast, high migration (1 × 10–2) resulted in catastrophic failure, reducing average CI coverage to just 22% (Figure 2a). This dramatic threshold effect indicates that the migration impact threshold lies between rates of 1 × 10−7 and 1 × 10–2.

Migration primarily compromises inferential reliability rather than precision alone. While root mean squared error (RMSE) increased modestly under high migration—roughly doubling from 0.00055 (no migration) to 0.00116 (high migration)—the collapse in CI coverage was far more severe. This 67 percentage-point drop in coverage (from 89% to 22%) indicates that migration introduces systematic bias in parameter estimates rather than merely increasing uncertainty around otherwise accurate estimates. Well-calibrated Bayesian inference should maintain nominal coverage even with increased uncertainty (i.e., wider credible intervals), but the near-zero coverage under high migration reveals fundamental model misspecification.

### Model misspecification bias, not increased uncertainty

Migration causes systematic bias where true parameters fall outside credible intervals, not increased uncertainty with maintained calibration. Across all 15 conditions with high migration (three patterns × five divergence times), the true divergence time failed to fall within the 95% CI in 100% of replicates (coverage = 0%). This complete systematic failure is qualitatively different from increased parameter uncertainty, which would manifest as maintained coverage probability with wider credible intervals. The pattern indicates that the Ecoevolity model, which assumes no post-divergence gene flow, systematically estimates incorrect divergence parameters when migration is present.

The divergence pattern complexity had minimal effect on this bias. For example, simultaneous divergence, grouped divergences, and independent divergences all showed 0% coverage under high migration. However, pattern complexity did affect precision in the absence of migration, with RMSE increasing from simple to complex patterns (0.00037 for simultaneous, 0.00055 for grouped, 0.00094 for independent divergences).

### Migration systematically biases model selection toward simpler scenarios

High migration leads to systematic underestimation of divergence event complexity, not merely reduced statistical power. Analysis of the posterior probabilities for different numbers of divergence events (Figures 1b, 2b and 3b) reveals that migration causes strong bias toward inferring fewer divergence events than occurred. Under high migration, single-event models received high posterior support even when the true model involved multiple independent divergences (Figure 3b, bottom row). This effect intensified with divergence age: at 16*N_e_* generations with high migration, the probability mass concentrated almost entirely on 1-2 event models regardless of the true complexity (Figure 3b, rightmost panels).

Importantly, this represents systematic model selection bias rather than simply reduced power to distinguish between models. Under low or no migration, the method correctly identified model complexity with high posterior probability, placing mass on 1-event models for truly simultaneous divergences (Top panels for Figures 1b, 2b and 3b) and distributing mass across 2-4 events for more complex scenarios (Middle and bottom panels for Figures 1b, 2b and 3b), top rows). High migration fundamentally alters these inferences, systematically favoring overly simple evolutionary scenarios.

### Joint effects and interactions

Divergence age and migration interact to exacerbate inference problems. While recent divergences (1*N_e_*–2*N_e_*) showed some resilience to migration effects—maintaining partial CI coverage and reasonable model selection—older divergences (8*N_e_*–16*N_e_*) failed completely under high migration. This age-dependent effect suggests that gene flow accumulating over longer periods since divergence increasingly confounds the evolutionary signal, making it progressively harder to distinguish the original divergence pattern from ongoing migration. The three-way interaction between migration rate, divergence pattern, and divergence age explained 94% of variance in CI coverage and 87% of variance in RMSE. However, migration rate alone accounted for the majority of explained variance (71% for coverage, 52% for RMSE), with pattern and age contributing more modest independent effects.

### Conclusion

Our results demonstrate that Ecoevolity’s full-likelihood Bayesian approach, despite its theoretical advantages over ABC methods, is not robust to post-divergence migration. Researchers applying this method must either: (1) verify through independent evidence that migration is negligible (less than approximately 1 × 10−7), (2) explicitly incorporate migration into the model, or (3) interpret results with extreme caution when migration is suspected. The systematic nature of the bias—consistently underestimating both divergence times and model complexity—means that standard model diagnostics may fail to detect the problem, as the method will appear confident in its incorrect inferences.

## Notes

### Competing Interest Statement

The authors have declared no competing interest.

